# Structural basis for the interaction between the bacterial cell division proteins FtsZ and ZapA

**DOI:** 10.1101/2024.07.18.604045

**Authors:** Junso Fujita, Kota Hibino, Gota Kagoshima, Natsuki Kamimura, Yuki Kato, Ryo Uehara, Keiichi Namba, Takayuki Uchihashi, Hiroyoshi Matsumura

## Abstract

Cell division in most bacteria is regulated by the tubulin homolog FtsZ as well as ZapA, a FtsZ-associated protein. However, how FtsZ and ZapA function coordinately has remained elusive. Here we report the cryo-electron microscopy structure of the ZapA-FtsZ complex. The complex forms an asymmetric ladder-like structure, in which the double antiparallel FtsZ protofilament on one side and a single protofilament on the other side are tethered by ZapA tetramers. In the complex, the extensive interactions of FtsZ with ZapA cause a structural change of the FtsZ protofilament, and the formation of the double FtsZ protofilament increases electrostatic repulsion. High-speed atomic force microscopy analysis revealed cooperative interactions of ZapA with FtsZ at a molecular level. Our findings not only provide a structural basis for the interaction between FtsZ and ZapA but also shed light on how ZapA binds to FtsZ protofilaments without disturbing FtsZ dynamics to promote cell division.

## Introduction

Cell division in nearly all bacteria is initiated by the polymerization of the tubulin homolog FtsZ at midcell^1–3^. In the presence of GTP, FtsZ polymerizes into protofilaments, which further associate into a ring-like structure (the Z-ring). The Z-ring has two important functions: recruiting the cell division proteins as a scaffold, and treadmilling of FtsZ protofilaments dependent on its GTPase activity. Treadmilling is a motion in which the FtsZ molecule binds to the protofilament’s end (plus-end) and depolymerizes at the other end (minus-end). FtsZ treadmilling is critical to promote condensation of diffuse FtsZ protofilaments into a coherent Z-ring, and also for stimulating septal cell wall synthesis, which is essential for cell constriction^4, 5^.

FtsZ-associated proteins (Zaps) such as ZapA, ZapB, ZapC, and ZapD facilitate Z-ring assembly^6–9^. *E. coli* mutants lacking one of Zap genes typically exhibit elongated cells, and *E. coli* mutants lacking multiple Zap genes show mislocalized and distorted Z-rings, resulting in a more severe phenotype and/or death. The best characterized Zap is ZapA, which is conserved widely in gram-negative and gram-positive bacterial species^6, 10, 11^. ZapA is usually expressed at a very high level in bacterial cells; for example, the cellular ZapA concentration in *E. coli* is estimated to be 5.4 μM, approximately equal to that of FtsZ^11^. FtsZ can assemble into helical or straight protofilaments *in vivo*^12–14^ and *in vitro*^15^. ZapA binds FtsZ straight protofilaments preferentially to crosslink FtsZ protofilaments^16^; thereby, ZapA bundles more aligned and straight filaments constituting a coherent Z-ring^11, 17^. A recent *in vitro* reconstituted FtsZ and ZapA study has revealed that ZapA binds to FtsZ protofilaments in a highly cooperative manner, and has no effect on treadmilling velocity^17^. Such interaction between FtsZ and ZapA is suggested to be important in maintaining a coherent and dynamic Z-ring. Thus, along with FtsZ, the importance of the ZapA function in bacterial cell division has gained attention.

Structures of FtsZ and ZapA from several bacterial species have been characterized independently. Recently, we have reported the high-resolution cryo-electron microscopy (cryo-EM) structure of *K. pneumoniae* FtsZ protofilaments, demonstrating the structural details of straight and helical protofilaments^15^. Crystallographic analyses of ZapA from *P. aeruginosa* and *E. coli* have been performed, showing that ZapA forms a dumbbell-like tetramer with domains at both ends that bind to FtsZ^10, 18^. The ladder-like structure of the *E. coli* ZapA-FtsZ complex has been previously observed by negative staining electron microscopy^11, 19^. Despite extensive studies, our understanding of ZapA-FtsZ coordination has been hampered by a lack of a high-resolution structure of the ZapA-FtsZ complex.

In this study, we report the high-resolution cryo-EM structure of the *K. pneumoniae* ZapA-FtsZ complex. The cryo-EM structure of the ZapA-FtsZ complex reveals an asymmetric ladder-like structure, in which the double antiparallel FtsZ protofilament on one side and a single protofilament on the other side are tethered by ZapA tetramers. The extensive interactions of FtsZ with ZapA cause a structural change in the FtsZ protofilament, shedding light on how ZapA cooperatively binds to FtsZ protofilaments. The electrostatic repulsion is found in the inter-filament interactions within the FtsZ double protofilaments, possibly explaining why ZapA binding does not affect FtsZ treadmilling velocity. We also performed high-speed atomic force microscopy (HS-AFM) analysis to investigate their single-molecule dynamics. The HS-AFM analysis indicates cooperative interactions of ZapA with FtsZ protofilaments at a molecular level. Taken together, our data provide a structural basis for the interaction between FtsZ and ZapA and lead to a mechanistic model for the regulation of bacterial cell division.

## Results

### The ZapA tetramer binds to antiparallel double protofilaments of FtsZ

We first observed the *K. pneumoniae* ZapA-FtsZ complex by negative staining EM and found many twisted ladder-like structures (Extended Data Fig. 1a), as observed previously^11, 19^. To determine the high-resolution structure, we performed cryo-EM data collection of the ZapA-FtsZ complex. Many paired filaments with aligned high-contrast dots were observed (Fig. 1a), and 2D class averages indicated the ladder-like structure (Fig. 1b). During picking by filament tracer, it quite often happened that one side of the filament in the ladder was centered instead of the middle of the ladder with a lower signal. Therefore, the reconstructed map showed clear features on one side of the ladder, but blurred features on the opposite side (Extended Data Fig. 1b). We therefore re-centered the initial 3D map and re-extracted particles for another reconstruction to observe the filaments on both sides of the ladder. The reconstructed map showed an asymmetric ladder structure; the FtsZ formed a double protofilament on one side, and a single protofilament on the other side, and these were tethered by ZapA tetramers (Extended Data Fig. 2a,b). The axes along the two FtsZ protofilaments were not completely parallel. Even after many trials, we failed to improve the resolution of the FtsZ single protofilament region, and therefore the polarity and orientation of the FtsZ single protofilament could not be determined. Hereafter, we focused on the FtsZ double protofilament bound to ZapA tetramer. We subtracted the single protofilament region and performed symmetry expansion with C2 symmetry and 3D classification to align “upside-down” double protofilaments. After removing the duplicated particles, we performed helix refine with D1 symmetry, namely with a single C2 axis perpendicular to the helical axis but no rotational symmetry along the helical axis^20^, and reached an overall map resolution of 2.73 Å (Extended Data Fig. 3a–c) with an optimized helical rise of 44.58 Å and a helical twist of −3.11° (Extended Data Table 1). The well-resolved FtsZ double protofilament structure enabled us to confirm that all FtsZ monomers are in the T conformation with GMPCPP bound and that the two FtsZ protofilaments are antiparallel (Fig. 1c,d). Secondary structural elements are mostly conserved in ZapA-free and ZapA-bound FtsZ (Extended Data Fig. 5a). One dimer head of ZapA was located among four FtsZ molecules. The dimer head of ZapA on the other side could not be modeled.

**Fig. 1:**
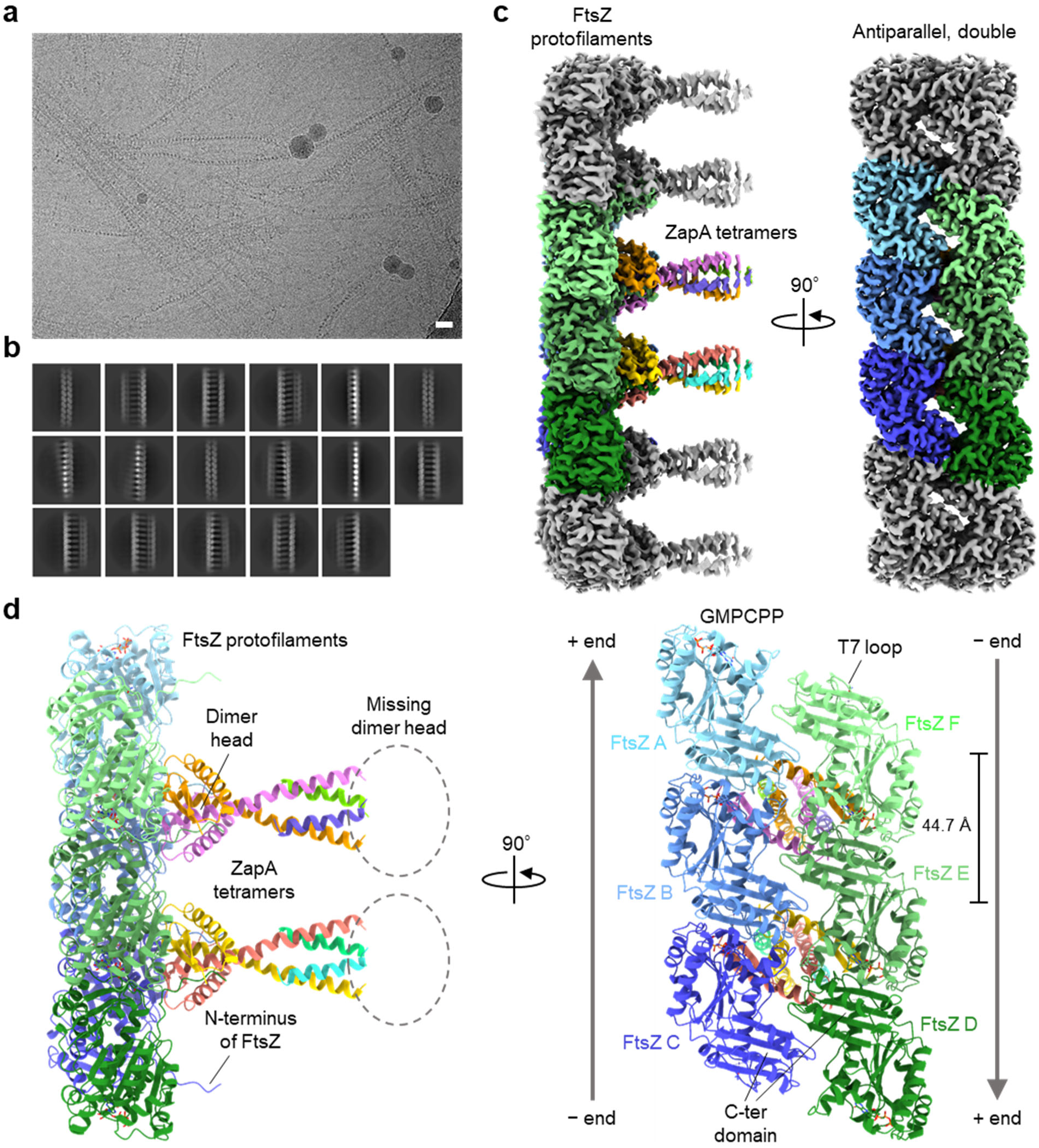
Overall cryo-EM structure of an antiparallel double filament of KpFtsZ in complex with KpZapA tetramer. **a**, Typical raw micrograph of the ZapA-FtsZ complex supplemented with 1 mM GMPCPP. The scale bar represents 20 nm. **b**, 2D class averages selected before ab-initio reconstruction. **c**, Final sharpened map at an overall resolution of 2.73 Å. The colored region corresponds to the region we reconstructed the model, and each color represents different chains. **d**, The reconstructed model containing six FtsZ monomers and two ZapA tetramers. The coloring is the same as in **c**. The directions of two antiparallel FtsZ protofilaments are shown in the right panel.

### Two ZapA molecules interact with four FtsZ molecules through the N-terminal tail of FtsZ

In the ZapA-FtsZ interface, the ZapA dimer head (chains G, H) interacted with two pairs of FtsZ dimers from two antiparallel protofilaments (chains A, B, and E, F) (Fig. 1d). As the ZapA-FtsZ complex has a C2 axis perpendicular to the protofilament, we focused on the interactions around one pair of FtsZ dimers (chains A, B). The N-terminal region of FtsZ (residues 1–10) was sandwiched between two ZapA molecules (Fig. 2a). The region from Leu41 to Tyr63 in one ZapA (chain G) mainly interacted with the N-terminal tail from Met1 to Thr8 in FtsZ (chain A) (Extended Data Fig. 4a). Thr45, Arg46, Val47, Thr48, and Asn60 in ZapA (chain G) formed an elaborate network of hydrogen-bonding interactions with Phe2, Met5, Glu6, and Leu7 in FtsZ (chain A). The region from Arg16 to Gln23 in the other ZapA (chain H) primarily interacted with the same N-terminal tail of FtsZ (chain A). Hydrogen bonds were observed between Gln28 in ZapA (chain H) and Met1 in FtsZ (chain A), and between Asn18 in ZapA (chain H) and Glu3 of FtsZ (chain A). The region from Asp7 to Arg16 in the same ZapA (chain H) interacted with the region from Arg33 to Gly55 in the other FtsZ (chain B) (Extended Data Fig. 4b). Gly12, Ser14, and Arg16 in ZapA (chain H) formed hydrogen-bonding interactions with Arg33, Thr52, Ala53, Val54, and Gly55 in FtsZ (chain B). The side chain of Asn49 in ZapA (chain G) made a hydrogen bond with the carbonyl oxygen of Lys51 in FtsZ (chain B).

**Fig. 2:**
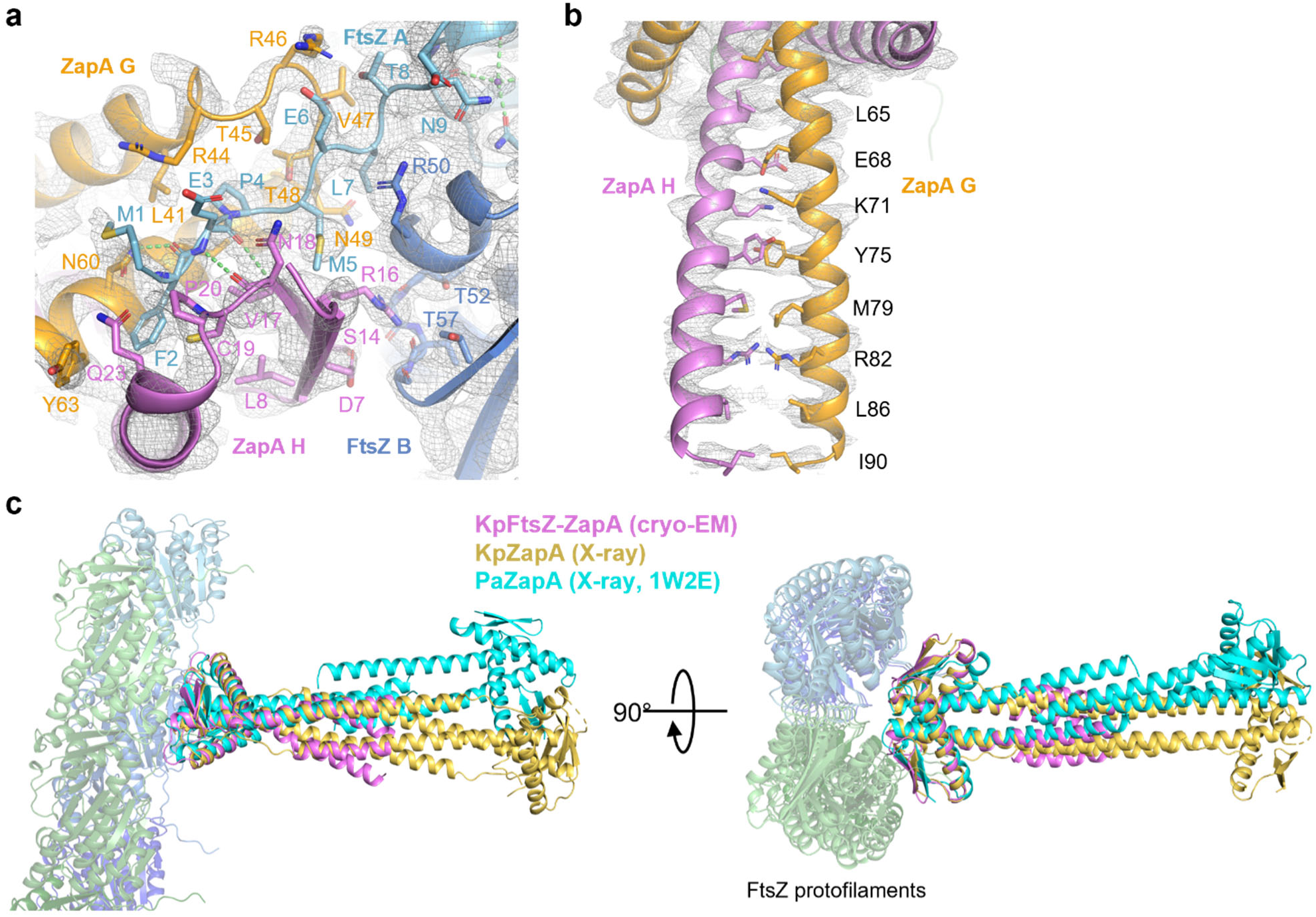
Interactions between FtsZ and ZapA. **a**, Interface between FtsZ and ZapA. N-terminus of FtsZ is located between two ZapA monomers. Final sharpened map is represented as a gray mesh. Hydrogen bonds are shown as green dashes. **b**, Bundled helices within one ZapA dimer. Another ZapA dimer is not shown for clarity. **c**, Superimposition of cryo-EM structure of the ZapA-FtsZ complex (violet) and crystal structures of KpZapA (yellow, this study) and PaZapA (cyan, PDB code: 1W2E). The structures are superimposed based on ZapA dimers.

Protein-protein interactions were analyzed using the PDBe PISA server^21^ (Extended Data Table 2). The interface area of the ZapA dimer head (chains G, H) with two pairs of FtsZ dimers is extensive (2,105.4 Å^2^) with a Δ*G* of dissociation of −17.0 kcal/mol, as compared with the interface area and a Δ*G* of dissociation between two FtsZ molecules within a FtsZ protofilament, which are 1,390.6 Å^2^ and −16.0 kcal/mol, respectively.

ZapA tetramers were stabilized by bundling of four C-terminal helices. In our structure, up to Ile90 was resolved in the ZapA dimer bound to the double protofilaments of FtsZ (Fig. 2b). In the other ZapA dimer, only residues 85–103 could be built, and the dimer head was missing due to weak density (Fig. 1c,d). To acquire the structural information of the missing dimer head, we determined the crystal structure of *K. pneumoniae* ZapA (KpZapA) at 1.8 Å resolution (Fig. 2c, Extended Data Table 3, Extended Data Fig. 5b) and performed a structural comparison of ZapA within the ZapA-FtsZ complex with the crystal structures of KpZapA, *E. coli* ZapA (EcZapA, PDB code: 4P1M^18^), and *P. aeruginosa* ZapA (PaZapA, PDB code: 1W2E^10^). The cryo-EM structure of KpZapA within the ZapA-FtsZ complex was superimposed well to the crystal structures of KpZapA and EcZapA (r.m.s.d. = 0.851 Å among 128 C_α_ atom for KpZapA, and 0.911 Å among 127 C_α_ atom for EcZapA) (Fig. 2c). For the dimer head on the missing side, KpZapA and EcZapA overlapped well but PaZapA did not. The N-terminal tail of FtsZ in our cryo-EM structure, which is sandwiched by two ZapA molecules (Fig. 2a), overlaps with the C-terminal tail of KpZapA and EcZapA (Extended Data Fig. 6a). This observation indicates that the C-terminal tail of ZapA moves aside to accept the N-terminal tail of FtsZ when the ZapA-FtsZ complex forms. Furthermore, the α1-α2 loop of ZapA undergoes a conformational change to recognize the N-terminal tail of FtsZ.

### Intra- and inter-FtsZ protofilament interactions

We found GMPCPP in the binding pocket between two FtsZ molecules in the protofilament, and the large density around the γ-phosphate enabled us to model two conformations of GMPCPP (Fig. 3a). In conformation 1, the γ-phosphate was located near the T3 loop, and the similar conformations have been observed in the previous crystal structures of FtsZ complexed with GTP or its analog (PDB code: 3WGN^22^) (Fig. 3b). The position of γ-phosphate in conformation 2 was close to that of our previous cryo-EM structure of the KpFtsZ single protofilament (PDB code: 8IBN^15^) (Fig. 3c). The distance between the γ-phosphate in conformation 2 and Asp209 in the upper FtsZ was 4.7 Å, which is closer than that between the γ-phosphate in conformation 1 and Asp212 of FtsZ (5.7 Å) (Fig. 3d). The H6-H7 loop and the C-terminal domain mainly contributed to the interaction between the two antiparallel protofilaments of FtsZ (Fig. 3e).

**Fig. 3:**
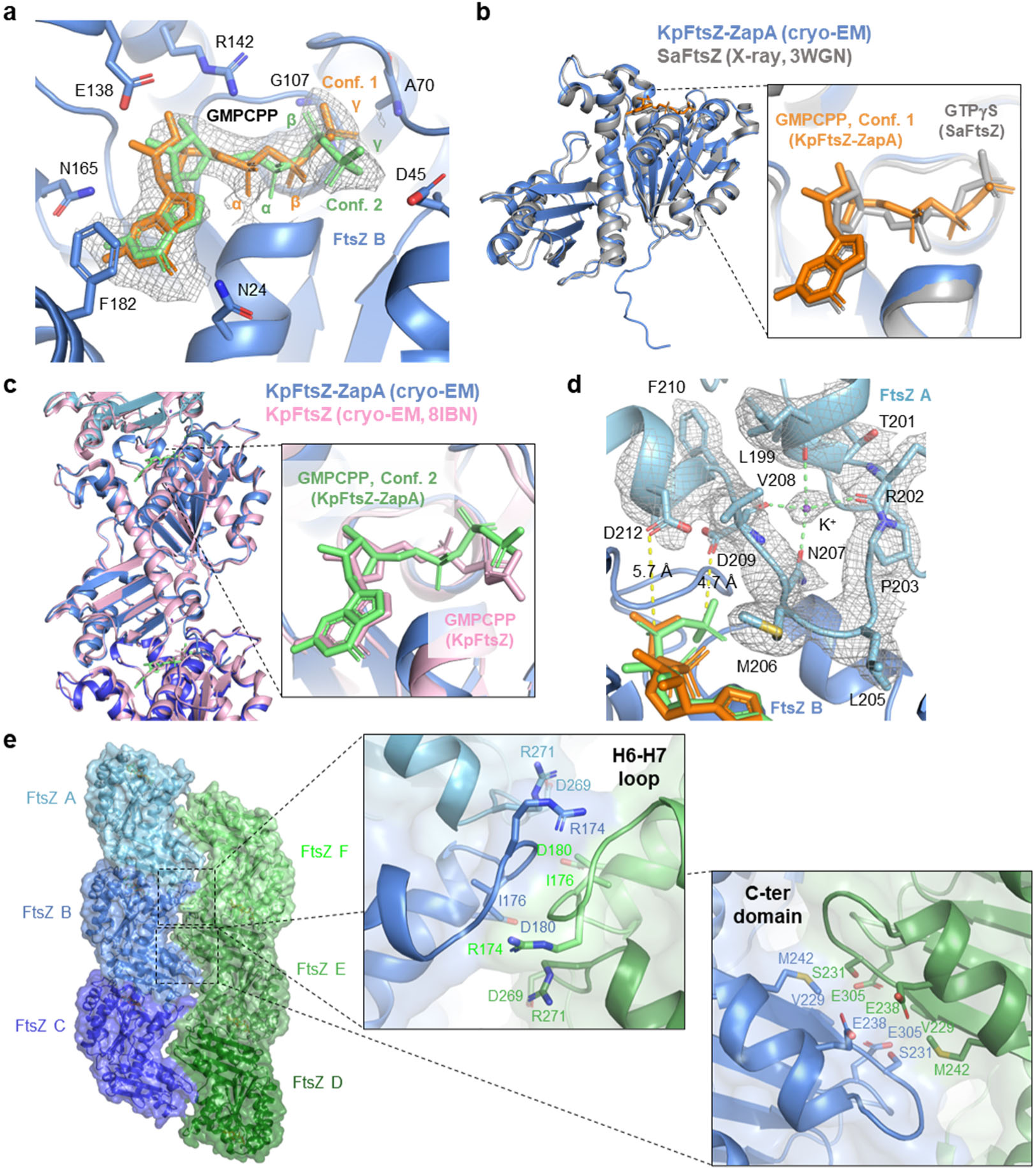
Interactions in FtsZ protofilaments. **a**, Close-up view around GMPCPP bound to FtsZ. Two alternative conformers are shown in orange and lime green. The final sharpened map is shown as a gray mesh and drawn only around GMPCPP. **b**,**c**, Superimposition of cryo-EM structure of the ZapA-FtsZ complex with GMPCPP (blue) and crystal structure of SaFtsZ with GTPγS (gray, PDB code: 3WGN, **b**) or cryo-EM structure of KpFtsZ with GMPCPP (pink, PDB code: 8IBN, **c**). Right panels are close-up views around GTP analogs. Only conformer 1 and 2 are shown in **a** and **b**, respectively. **d**, Close-up view around the T7 loop of FtsZ with potassium ion bound. The map is drawn only around the T7 loop. **e**, Interfaces between two antiparallel FtsZ protofilaments. Surface of FtsZ protofilaments are also shown. Right panels are close-up views around the H6-H7 loop and C-terminal domain. ZapA is not shown for clarity.

Inter-protofilament interactions within the double FtsZ filaments appear to be weak. According to the PISA calculation, chain B and chains F and E form an interaction area of 523.4 Å^2^ with a Δ*G* of dissociation of −3.6 kcal/mol (Extended Data Table 2). This feature is rationalized by the electrostatic repulsions among Glu238 and Glu305 found in the inter-protofilament interactions (Fig. 3e).

The entire structure of the FtsZ protofilament with ZapA was roughly similar to that without ZapA (r.m.s.d. = 1.529 Å among 918 C_α_ atoms for three KpFtsZ molecules within a protofilament, Fig. 3c), but ZapA binding induces structural changes in a FtsZ molecule as well as an adjacent FtsZ molecule. When comparing the structures of FtsZ within the ZapA-FtsZ complex and that of FtsZ straight protofilaments alone (PDB code: 8IBN^15^), approximately 2−3 Å shifts of the α1-β2 loop and the α2-β3 loop are observed (indicated by red and green arrows in Fig. 4). The β-sheet comprising β-strands 1-5 slightly shifts upon ZapA binding (indicated by a blue arrow in Fig. 4). The side chains of Asp209 and Phe210 in FtsZ also rearrange. This conformational change allows Asp209 to form a hydrogen bond with the γ-phosphate oxygen of the GMPCPP in conformation 2.

**Fig. 4:**
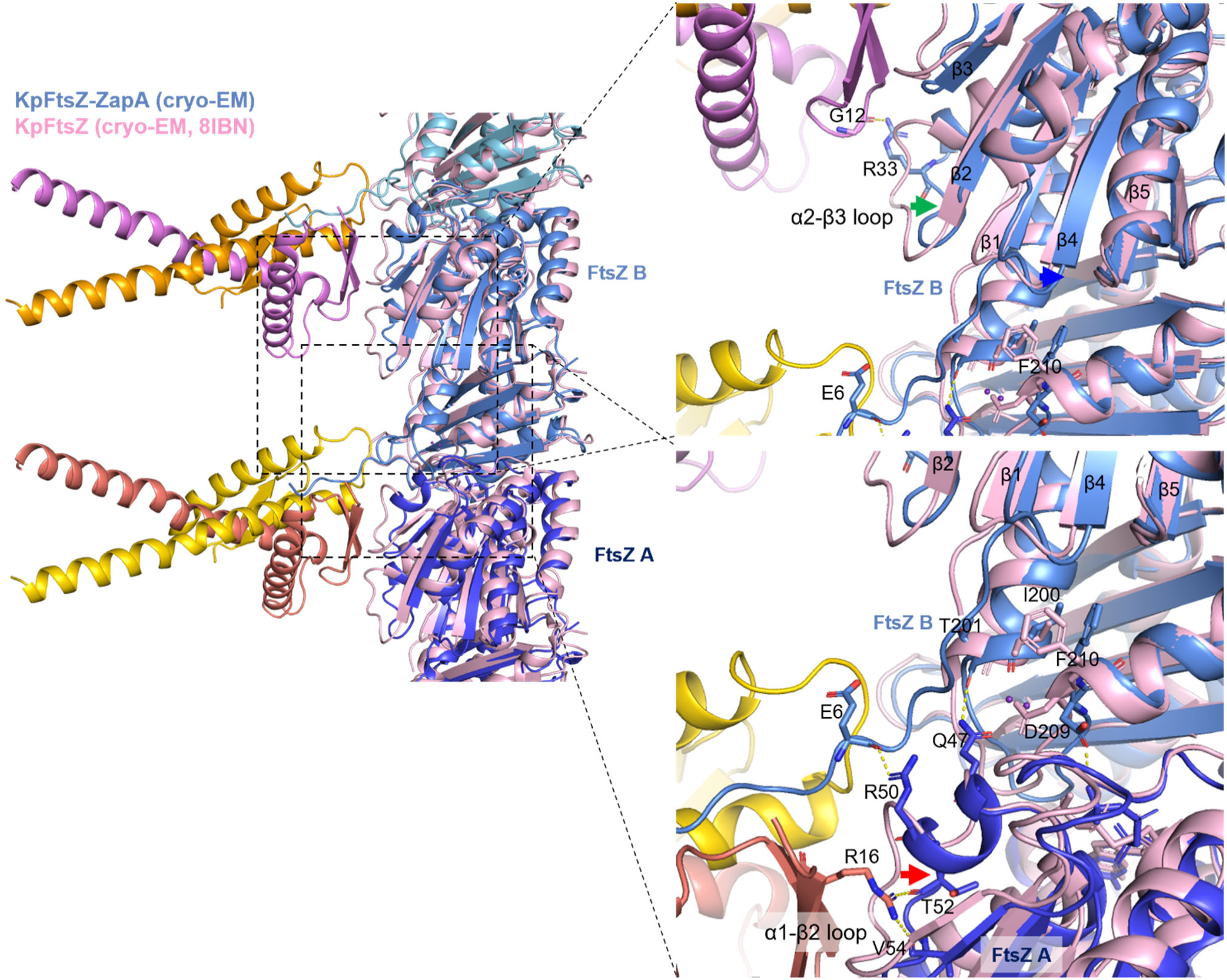
Superimposition of the ZapA-FtsZ complex and the KpFtsZ straight protofilaments alone. The coloring of the KpFtsZ-ZapA complex is the same as in Fig. 1**d**, and the KpFtsZ straight protofilament alone is colored in pink. Several key residues at the KpFtsZ-ZapA interface are shown as sticks and labeled. The red, blue, and green arrows indicate the structural shifts or changes of α1-β2 1oop, β-sheet comprising of β1-5, and α1-β1oop, respectively.

### HS-AFM analysis of the ZapA-FtsZ complex

To investigate the single-molecule dynamics of the ZapA-FtsZ complex, we performed HS-AFM analysis. We first observed the ZapA-FtsZ complex on a mica substrate after mixing FtsZ, ZapA, and GTP in a tube, and it revealed ladder-like structures in which ZapA molecules tether to the FtsZ protofilaments (Extended Data Fig. 7a). Judging from the measured heights from the cross-sectional profile (Extended Data Fig. 7a), ZapA appeared to bridge two single FtsZ protofilaments on the mica substrate to form the ladder-like structures. As far as we observed by HS-AFM, the structures of ZapA bridging double FtsZ protofilaments and single protofilaments, as resolved by cryo-EM, were not observed. Since the longitudinal axes of the double protofilament and single protofilament cross-linked by ZapA are not parallel according to the cryo-EM analysis, there is a possibility that structural distortion upon adsorption onto the planar substrate prevents the maintenance of the native complex structures. Therefore, the ladder structures observed by HS-AFM might be artifacts resulting from the disruption of ZapA-mediated cross-linking between double protofilaments and single protofilaments, converting them into cross-linking of single protofilaments. This might indicate that the interaction between the double protofilaments and ZapA is weak and easily disrupted.

Next, we pre-adsorbed FtsZ protofilaments onto a mica substrate (Fig. 5a) and then added ZapA (final concentration: 0.6 μM) to the solution. Figs. 5a and b show HS-AFM images before and after the addition of ZapA, respectively. When ZapA was added, bright spots were clearly observed on the FtsZ protofilaments (Fig. 5b). A magnified view shows that ZapA binds to the hollow sites between two FtsZ protofilaments as well as on top of a single FtsZ protofilament (Fig. 5c). This is considered to correspond to the binding of ZapA to FtsZ double protofilaments and single protofilaments revealed by cryo-EM. This suggests that the adsorption surface of FtsZ to the mica substrate is opposite to the ZapA binding surface, which may be why we cannot directly observe ZapA bridging between double protofilaments and single protofilaments.

**Fig. 5:**
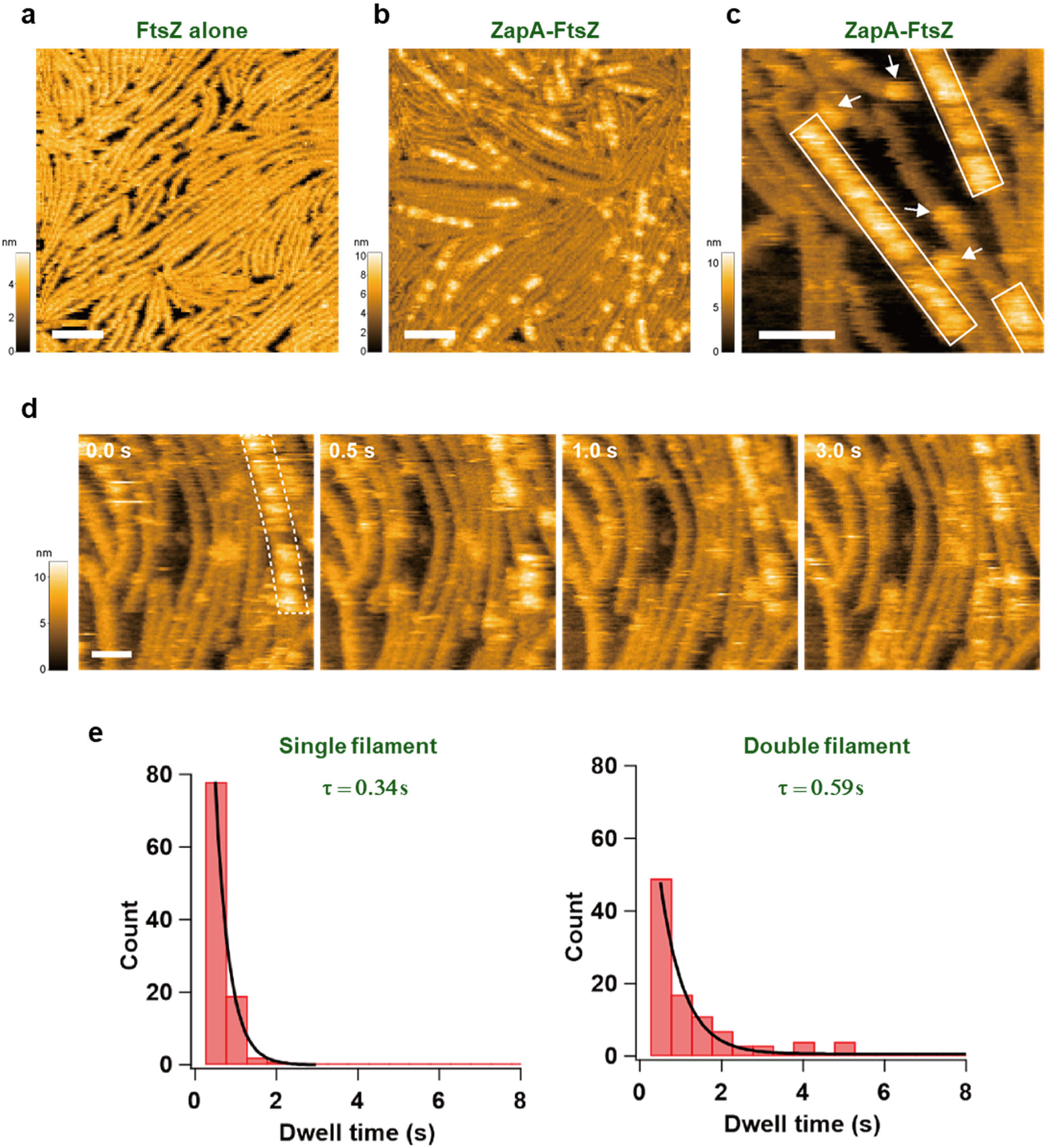
Interaction between FtsZ Filaments and ZapA Observed by HS-AFM. **a**, HS-AFM image of FtsZ filaments alone. Scale bar: 50 nm, frame rate: 1 frame per second (fps), 150 × 150 pixels. **b**, **c,** HS-AFM images of FtsZ filaments interacting with ZapA. Scale bar: 50 nm, frame rates: (b) 1 fps, (c) 2 fps, 150 × 150 pixels. Rectangles encircle regions where ZapA is bound to the FtsZ double filaments, and arrows indicate ZapA bound to a single FtsZ filament. **d**, Successive HS-AFM images showing the dynamic interaction between ZapA and an FtsZ filament. Scale bar: 30 nm, frame rate: 2 fps, 135 × 135 pixels. Regions where ZapA is bound to the double filaments are encircled by a broken line at 0 s. **e**, Histograms of the dwell time of ZapA bound to the double (left) and single (right) filaments (*n* = 100 for both). The solid lines show fitting curves with an exponential function.

The binding of ZapA to the FtsZ protofilaments is not static but dynamic, with repeated binding and dissociation (Fig. 5d and Supplementary Video 1). We analyzed the binding residence time of a single bright spot, which is likely corresponding to a ZapA tetramer, on FtsZ protofilaments. The residence time on single protofilaments was 0.34 s, while on double protofilaments it was 0.57 s, about 1.7 times longer, indicating that ZapA has a higher affinity for double protofilaments than single protofilaments. It should be noted that this residence time analysis was performed on isolated bright spots binding to protofilaments. As seen in Fig. 5d, there are areas where ZapA bright spots are contiguous along a protofilament, especially when bound to double protofilaments. This indicates that when ZapA molecules are adjacent to each other, their affinity for FtsZ protofilaments increases, demonstrating positive cooperativity in binding.

## Discussion

Because both FtsZ and ZapA are widely conserved among bacteria and an interplay between FtsZ and ZapA is pivotal for bacterial cell division, the structural information of the FtsZ-ZapA complex is essential to understand the mechanism of bacterial cell division. The cellular ZapA concentration of in *E. coli* is approximately 5.4 μM, roughly equal to that of FtsZ^11^. The number of FtsZ molecules in *E. coli* is estimated to be between 3,200^23^ and 15,000^24^. Consequently, during the initial stage of cytokinesis, FtsZ self-polymerizes and associates into condensed protofilament bundles to form the Z-ring, and ZapA molecules bind to any part of the Z-ring to stabilize it. A FtsZ protofilament has polarity and lacks symmetry, while ZapA is in an equilibrium between homodimer and homotetramer, which are related by C2 and D2 symmetry, respectively. How do the symmetric ZapA molecules bundle the asymmetric FtsZ protofilaments at multi-point interactions? We have determined the FtsZ-ZapA complex at high resolution to answer this question. Our cryo-EM analysis shows that ZapA dimer bundles the double antiparallel FtsZ protofilament. Moreover, our cryo-EM and HS-AFM analyses show at least two interaction modes between ZapA and FtsZ, in which ZapA interacts with double antiparallel protofilaments as well as a single protofilament. The different interaction modes can be explained by a combination of ZapA tetramer symmetry and multi-point interactions (Fig. 6a). If the FtsZ protofilaments on both sides were both double, the whole structure would have D2 symmetry because the ZapA tetramer also has D2 symmetry (Fig. 6a, left panel). However, based on the orientations of the two dimer heads of ZapA, such D2 symmetry would not be acceptable. Therefore, the interactions with a single protofilament on only one side will give an asymmetric structure (Fig. 6a, right panel). We assume that the asymmetric structure of the ZapA-FtsZ complex may help generate curvatures and dynamic behavior because the single protofilament should be more flexible than the double antiparallel protofilament. Considering that FtsZ protofilaments have to be moderately curved in the Z-ring, a double and single protofilament of FtsZ may be located on the outer and inner sides, respectively (Fig. 6b). As FtsA forms antiparallel double filaments in the presence of FtsN^28^, the antiparallel double filaments of FtsZ stabilized by ZapA may be tethered to those type of FtsA filament with 1:1 interaction.

**Fig. 6:**
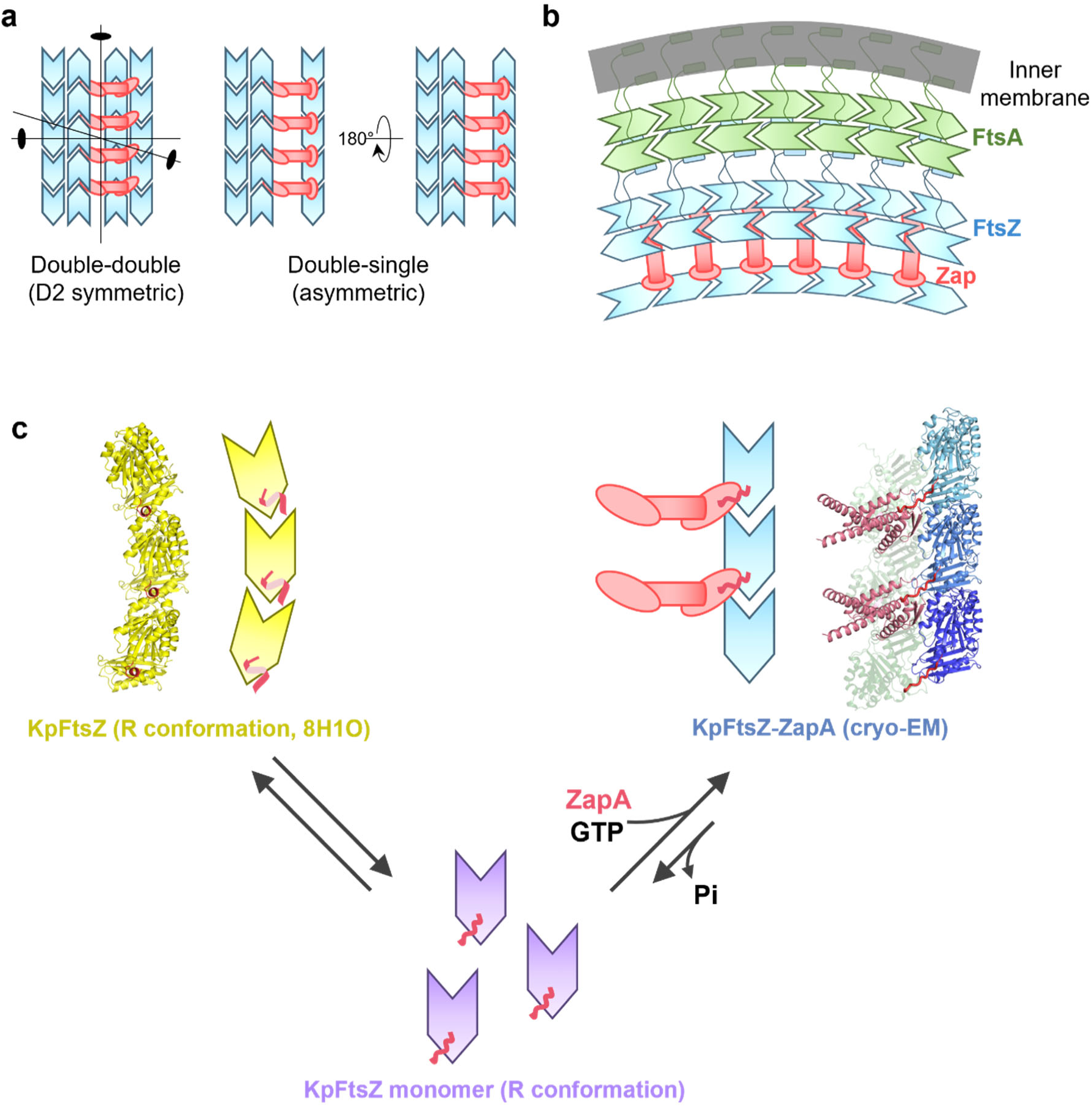
Assembly model of the ZapA-FtsZ. **a**, Three types of possible ZapA-FtsZ complex. If two heads of ZapA tetramer are symmetric, FtsZ double protofilaments on each side should be also symmetric, therefore the complex has D2 symmetry (left panel). Otherwise, double and single FtsZ protofilament are generated in each side with no symmetry. This asymmetric complex has one C2 axis if the direction of the single protofilament is ignored (right panel). **b**, Model of interactions between FtsA and the ZapA-FtsZ complex. Double antiparallel FtsZ protofilaments stabilized by ZapA tetramers can bind to double antiparallel FtsA filaments activated by FtsN through the flexible C-terminal linker. **c,** FtsZ conformational equilibrium model regulated by ZapA. The N-terminal tail of FtsZ (magenta) adopts a short helix in GDP-bound helical FtsZ protofilament, and the monomeric form of FtsZ must adopt a flexible N-terminal tail, while the tail is captured by ZapA to be extended in the ZapA-FtsZ complex.

Depending on the GTP/GDP ratio, FtsZ adopts monomer conformations as well as various assembly forms. The GTP-bound FtsZ adopt a straight protofilament form, while the GDP-bound FtsZ can be either monomers or curved protofilaments. During the early stage of cell division, GTP triggers FtsZ molecules to form straight protofilaments, and ZapA crosslinks the FtsZ protofilaments to bundle more aligned protofilaments^11, 17^, constituting a coherent Z-ring. This maturation of the Z-ring is promoted by ZapA’s property of preferentially binding to straight FtsZ protofilaments^16^. We consider that this behavior of ZapA may be facilitated by its interaction through the N-terminus of FtsZ. The monomeric form of FtsZ has a flexible N-terminal tail. In the helical (curved) protofilaments, the FtsZ’s N-terminal tail adopts a short helix to stabilize the curved conformation, revealed by our previous cryo-EM structure of the GDP-KpFtsZ complexed with a monobody^15^ (Extended Data Fig. 6b). We have now identified the direct interaction between ZapA and the N-terminal tail of FtsZ that forms straight protofilaments. Based on the finding, we propose a model of FtsZ conformational equilibrium regulated by ZapA (Fig. 6c). In this model, when FtsZ protofilaments are captured by ZapA molecules, the conformation equilibrium shifts toward FtsZ straight protofilament bundles. During treadmilling of FtsZ protofilaments, the released GDP-bound FtsZ molecules and remaining GDP in the cells still favor the monomeric or curved FtsZ protofilament conformation. However, the interactions with ZapA through FtsZ’s N-terminal tail may prevent such a backward shift of the equilibrium to stabilize the more aligned bundled protofilaments for the promotion of cell division. Recent studies have suggested a "cytomotive switch" model in which the conformational switch of FtsZ is induced by polymerization and depolymerization^25–27^, rather than GDP/GTP. In addition to the model, a hierarchical model in which interactions with FtsZ binding proteins such as ZapA contribute to biasing a FtsZ conformational equilibrium is conceivable.

We also discuss a mechanistic insight into the cooperative interactions between ZapA and FtsZ. In our HS-AFM analysis, cooperative binding of ZapA to FtsZ filaments was observed at the single-molecule level. This is consistent with the previous experiments by total internal reflection fluorescence microscopy (TIRFM) showing a high cooperativity with a Hill coefficient much greater than 1^17^. Our cryo-EM analysis of the ZapA-FtsZ complex shows no direct interactions between ZapA tetramers. Therefore, the cooperativity is likely caused by structural changes in FtsZ protofilaments. Indeed, we found such structural changes upon ZapA binding to make the adjacent FtsZ molecule more accessible for the next ZapA molecule binding. The interactions of Arg16 of ZapA with Thr52 and Val54 of FtsZ (chain A), those of Glu6 of FtsZ (chain B) and Arg50 of FtsZ (chain A), and those of Thr201 of FtsZ (chain B) and Gln47 of FtsZ (chain A) appear to cause an approximately 2−3 Å shift of α1-β2 loop (indicated by a red arrow in Fig. 4). Such interactions likely rearrange the side chains of Asp209 and Phe210 of FtsZ (chain A), thereby shifting β-sheet comprising of β1-5, as indicated by a blue arrow in Fig. 4. This shift, in turn, causes the shift of α2-β3 loop (residues 31−36) (indicated by a green arrow in Fig. 4), in which Arg33 in the adjacent FtsZ molecule interacts with Gly12 of ZapA We consider that such extensive interactions between ZapA and FtsZ are key to increase positive cooperativity.

Previous TIRFM experiments have suggested that ZapA binds to FtsZ protofilaments without affecting FtsZ treadmilling velocity^17^. The interaction between two FtsZ protofilaments increases electrostatic repulsion, leading to almost no gain in Δ*G* of dissociation (−3.6 kcal/mol for chain B and chains E and F) (Extended Data Table 2), which is considered essential to form a coherent Z-ring. The increased electrostatic repulsion between protofilaments may explain how closely-bundled FtsZ protofilaments can treadmill without interfering with each other.

## Online Methods

### Protein expression and purification

KpFtsZ was expressed, and purified as we described previously^15^. The KpZapA coding oligonucleotide was chemically synthesized (Integrated DNA Technologies), and cloned into modified pCold I vector^15^ (Takara Bio) using the In-Fusion HD Cloning kit (Takara Bio) and primers KpZapA_forward: 5’-TCTAGATAGGTAATCTCTGCTTAAAAG-3’ and KpZapA_reverse: 5’-CATATGACCCTGGAAATAAAGATTCTC -3’. The *E. coli* cells overexpressed KpZapA with N-terminal to His6 tag and a TEV cleavage site were harvested at 4 °C, washed once with buffer A (50 mM Tris-HCl pH 7.5, 300 mM NaCl), and broken by ultrasonication on ice. After ultracentrifugation, the soluble fraction was applied to a 5 ml HisTrap HP column (Cytiva). Elution was carried out with a gradient of eluted with 45–500 mM imidazole in buffer A. Peak fractions were collected, and His6 tag was removed with 0.2 μM His-tagged TEV protease while dialyzing overnight against buffer A. His6 tag and His-tagged TEV protease were removed by passing through a second 5 ml HisTrap HP column. Protein fractions were pooled and concentrated on a Vivaspin 20 (MWCO; 5,000, Sartorius) to 4 ml and loaded onto a HiLoad 16/600 Superdex200 size-exclusion column (120 ml, Cytiva) equilibrated in buffer B (20 mM HEPES-NaOH pH 7.5, 150 mM NaCl).

### Negative staining

Carbon sides of amorphous carbon grids were glow discharged by using a JEC-3000FC sputter coater (JEOL). 3 μl of purified ZapA-FtsZ complex at a concentration of 1.0 mg ml^−1^ was loaded on each grid and blotted. Then each grid was immediately stained with 3 μl of 2% uranyl acetate solution and blotted, and this process was repeated three times. Each grid was air-dried for 30 min. Images were taken using JEM-1400Flash (JEOL, Japan) operated at 100 kV.

### Cryo-EM specimen preparation and data collection

The solution containing 12.3 μM (0.5 mg ml^−1^) KpFtsZ, 25.0 μM (0.32 mg ml^−1^) KpZapA, 1 mM GMPCPP in the buffer containing 16 mM HEPES pH 7.5, 14 mM NaCl, 100 mM KCl, and 5 mM MgCl2 were firstly prepared. This solution was diluted with the same buffer to a final concentration of the solution containing 7.4 μM (0.3 mg ml^−1^) KpFtsZ, 14.9 μM (0.19 mg ml^−1^) KpZapA (2× molar excess to FtsZ), 0.6 mM GMPCPP, 17 mM HEPES pH 7.5, 8.6 mM NaCl, 60 mM KCl, and 3 mM MgCl2, and the diluted sample solution was incubated on ice for 15 min before the freezing. Quantifoil grids (R1.2/1.3 Cu 200 mesh) were glow-discharged using a JEC-3000FC sputter coater (JEOL, Japan) at 20 mA for 20 seconds. 3 μl of the diluted sample solution was applied to the glow-discharged grids in a Vitrobot Mark IV chamber (Thermo Fisher Scientific, USA) equilibrated at 4 °C and 100% humidity. The grids were blotted with a force of –10 and a time of 1.5 sec and then immediately plunged into liquid ethane. Excess ethane was removed with filter paper, and the grids were stored in liquid nitrogen. Cryo-EM image datasets were acquired using SerialEM ver. 4.0^28^, yoneoLocr ver. 1.0^29^, and JEM-Z300FSC (CRYO ARM^TM^ 300: JEOL, Japan) operated at 300 kV with a K3 direct electron detector (Gatan, Inc.) in CDS mode. The Ω-type in-column energy filter was operated with a slit width of 20 eV for zero-loss imaging. The nominal magnification was 60,000×, corresponding to a pixel size of ∼0.87 Å. Defocus varied between −0.5 μm and −2.0 μm. Each movie was fractionated into 60 frames (0.038 s each, total exposure: 2.29 s) with a total dose of 60 e^−^/Å^2^.

### Cryo-EM image processing and model building

The gain reference was generated from 500 movies in the dataset with the "relion_estimate_gain" program in RELION 4.0^30^. The images were processed using cryoSPARC ver. 4.2.1^31^. 7,375 movies of the dataset were imported and motion corrected, and the contrast transfer functions (CTFs) were estimated. 6,070 micrographs whose CTF max resolutions were beyond 5 Å were selected. To prepare a 2D template, 35,635 particles were automatically picked from 500 micrographs using a filament tracer with the parameters; filament diameter, 200 Å; separation distance between segments, 0.5; minimum filament diameter, 150; maximum filament diameter, 220. The particles were extracted with a box size of 600 pixels with 2x binning. After two rounds of 2D classification, 2D class averages of ladder-like structures located near the center of the box were selected as templates. 4,374,348 particles were automatically picked from all micrographs using the templates in filament tracer with the parameters; filament diameter, 44 Å; separation distance between segments, 1; minimum filament length to consider, 5; angular sampling, 0.5°; standard deviation of gaussian blur, 0.4; hysteresis low threshold, 80; radius around crossings to ignore, 0.5. The particles were extracted with a box size of 600 pixels with 2x binning. After two rounds of 2D classification, 565,996 particles were selected and extracted with a box size of 480 pixels binning to 300 pixels.

The extracted particles were subjected to Ab-initio reconstruction with four classes. To improve the resolution of the FtsZ single protofilament region, we tried many things including re-centering of the map and local refinement, but all attempts were not successful. Therefore, hereafter we focused on the region of FtsZ double protofilament bound to ZapA tetramer. 148,091 particles were selected as a class showing FtsZ double protofilament and ZapA tetramer, and subjected to another Ab-initio reconstruction with a single class and maximum resolution of 6 Å. After symmetry search utility, helix refine was performed with an estimated helical rise of 44.67 Å and a helical twist of −1.965°. Particle subtraction was conducted to subtract the signals from other than FtsZ double protofilament and ZapA tetramer, and the subtracted particles were subjected to another helix refine with the reversed mask. The generated volume, mask, and particles were aligned with C2 symmetry using volume alignment tools, and C2 symmetry expansion was performed. The symmetry-expanded particles were classified into four classes with 3D classification, and the selected particles were subjected to the removal of duplicate particles. After the map of the best 3D class was aligned back to the original axis (helical axis corresponds to z-axis), helix refine, global and local CTF refine, and another helix refine were performed. Then the particles were subjected to local motion correction with an extracted box size of 360 pixels and a binning to 256 pixels, corresponding to a pixel size of 1.235 Å. The extracted particles were subjected to 2D classification, and 54,670 particles were selected. The selected particles were extracted with a box size of 320 pixels without binning and subjected to another round of helix refine with D1 symmetry, which means a single C2 axis perpendicular to the helical axis but no rotational symmetry along the helical axis^20^. After another round of global and local CTF refine and the final helix refine with D1 symmetry, the map resolution reached 2.73 Å (FSC=0.143) with the optimized helical rise of 44.58 Å and the helical twist of −3.11°. The entire workflow is shown in Extended Data Fig. 1b.

The model of the ZapA-FtsZ complex was built using the cryo-EM structure of KpFtsZ single protofilament (PDB code: 8IBN^15^) and the crystal structure of ZapA from *E*. *coli* (PDB code: 4P1M^18^) as initial models. After the initial models were manually fitted into the map using UCSF Chimera ver. 1.16^32^ and modified in Coot ver. 0.9.6^33^, real-space refinement was performed in PHENIX ver. 1.19.2^34^. The model was validated using MolProbity ver. 4.5.2^35^ in PHENIX ver. 1.19. 2^34^, and this cycle was repeated several times. The final model contains six FtsZ monomers (three monomers in a single protofilament) and two ZapA tetramers.

Protein-protein interactions were analyzed with PISA server^21^. Figures were prepared using ImageJ ver. 1.53q^36^, UCSF Chimera ver. 1.16^32^, ChimeraX ver. 1.6.1^37^, CLUSTALX ver. 2.0^38^, ESPript ver. 3.0^39^ (https://espript.ibcp.fr), LigPlot+ v.2.2^40^, and PyMOL ver. 2.5.0 (Schrödinger, LLC, USA).

### Protein crystallization, crystallographic data collection, processing, and refinement

Purified KpZapA at a concentration of 10 mg ml^−1^ was crystallized by hanging-drop vapor diffusion at 293 K (1 μl protein solution + 1 μl reservoir solution) with the reservoir solution consisting of 0.2 M Sodium acetate trihydrate, 0.1 M Sodium citrate pH5.5, and 10% PEG4000. Crystals were flash-cooled in a stream of nitrogen at 100 K without cryoprotectants after mounting in a loop. X-ray diffraction data were collected at a wavelength of 0.900 Å on the micro-focus beamline BL41XU at SPring-8, Hyogo, Japan using an EIGER X 16M detector (Dectris). The datasets were integrated and scaled using the KAMO system^41^ which runs BLEND^42^, XDS ver. 5 (February 2021)^43^, and XSCALE ver. 5 (February 2021)^43^ automatically. The phases for each structure were determined by molecular replacement with MOLREP in the CCP4 suite ver. 7.1^44^ using the previously determined structure of EcZapA (PDB code: 4P1M^18^) as the search model. Each model was refined with REFMAC ver. 5.8.0267^45^ and PHENIX ver. 1.19.2^34^, with manual modification using Coot ver. 0.8.6^33^. The refined structures were validated with MolProbity ver. 4.5.2^35^. Data-collection and refinement statistics are shown in Extended Data Table 3.

### HS-AFM measurement and data analysis

HS-AFM observations were carried out using a laboratory-built system. The operating mode of the high-speed AFM is the so-called tapping mode, in which the mechanical interaction between the needle probe and the sample is detected by vibrating the cantilever close to its resonant frequency and bringing it into intermittent contact with the sample^46^. The cantilever for the high-speed AFM is an Olympus AC10 with a resonant frequency in water of ∼500 kHz and a spring constant of ∼0.1 N/m. As no sharp probes were designed for the tip of the high-speed AFM cantilever, an amorphous carbon probe was fabricated by electron beam deposition and further sharpened by plasma etching in an Ar atmosphere to obtain a probe with a tip radius of curvature of less than about 5 nm^47^.

For the ZapA-FtsZ complex shown in Extended Data Fig. 7, following the same procedure as the sample subjected to cryo-EM analysis, 14 μM KpFtsZ, 36 μM KpZapA, and 1 mM GTP were mixed in a buffer solution (20 mM HEPES-NaOH pH 7.5, 5 mM MgCl2, 100 mM KCl), incubated for 5 minutes, and adsorbed onto a freshly cleaved clean mica substrate. After 10 minutes of incubation, it was washed with the same buffer to remove excess unadsorbed molecules. Subsequently, HS-AFM observation was performed in the buffer solution containing 1 mM GTP. For the data in Fig. 5, 8.8 μM KpFtsZ and 1 mM GMPPNP were mixed in the buffer solution to form protofilaments in advance, which were then adsorbed onto freshly cleaved mica and allowed to incubate for 5 minutes before washing. Then, the preformed FtsZ protofilaments were confirmed by HS-AFM in the buffer solution containing 1 mM GPMMPNP or 1 mM GMPCPP, after which ZapA was added to the observation solution to a final concentration of 0.6 μM to observe the interaction between FtsZ protofilaments and ZapA. All measurements were performed at room temperature.

The HS-AFM images shown in the manuscript were processed with tilt compensation and smoothing filters to reduce the image noise. The analysis of the dynamic interaction of ZapA on FtsZ protofilaments was performed by measuring the time from binding to dissociation for isolated ZapA bright spots on the images, as far as possible avoiding spots with neighboring molecules, and creating histograms based on this data. Image filtering and analysis of HS-AFM images were performed using custom image processing software developed with Igor Pro 9.0 (WaveMetrics Inc.,Lake Oswego, OR).

## Data availability

Cryo-EM atomic coordinates and maps of *K. pneumoniae* FtsZ double filament-ZapA tetramer have been deposited in the Protein Data Bank (PDB) and the Electron Microscopy Data Bank (EMDB) under the accession codes 9ISK and EMD-60837. Coordinates and structure factors of *K. pneumoniae* ZapA have been deposited in PDB under the accession number 9ISJ. The other coordinates used in this study are available from PDB. Source data are provided in this paper.

## Acknowledgments

We thank Yoshie Kushima and Reiko Yamauchi for their help in negative staining. This work was supported by: JSPS KAKENHI grant JP20K22630 (J.F.), JP23K06418 (R.U.), JP24K01994 (H.M.), JP24H02277 (H.M.), JP24H02270 (H.M.), JP23K18033(H.M.); Uehara Memorial Foundation (H.M.); Nagase Science and Technology Foundation (H.M.); Novartis Foundation (H.M.); G-7 Foundation (R.U.); JST OPERA (Open Innovation with Enterprises, Research Institute and Academia) grant JPMJOP1861 (K.N.); the Program for the R-GIRO Research from the Ritsumeikan Global Innovation Research Organization, Ritsumeikan University (H.M.); AMED BINDS (Platform Project for Supporting Drug Discovery and Life Science Research (BINDS) grant JP21am0101117 and JP22ama121003 (K.N.), and JP23ama121001 (H.M.); AMED CiCLE (Cyclic Innovation for Clinical Empowerment) grant JP17pc0101020 (K.N.); JEOL YOKOGUSHI Research Alliance Laboratories of Osaka University (K.N.); the Cooperative Research Program of the Institute for Protein Research, Osaka University (CR-22-02 and CR-23-02).

## Author contributions

J.F. and H.M. designed the research. K.H., N.K., Y.K., and R.U. prepared the protein sample and performed crystallization, crystallographic data collection, processing, refinement, and model building. J.F., K.H., and N.K. performed negative staining and cryo-EM data collection. J.F. performed cryo-EM image processing and model building. G.K. and T.U. performed HS-AFM experiments and analyzed the data. J.F., T.U., and H.M. prepared the figures and wrote the first draft of the manuscript. R.U. and K.N. helped to analyze and interpret the data and critically revise the manuscript. J.F. and H.M. conceptualized the study, developed the study design, supervised the authors throughout the study, and provided expertise in manuscript preparation. All authors read and approved the reviewed manuscript.

## Additional information

**Extended data** is available for this paper at https://doi.org/

## Supplementary information

The online version contains supplementary material available at https://doi.org/

