## Supplementary Information for "Structural basis for the interaction between the bacterial cell division proteins FtsZ and ZapA"

**This supplementary information contains:**
**Extended data Fig. 1–7**
**Extended data Table 1–3**
**Supplementary Video 1**

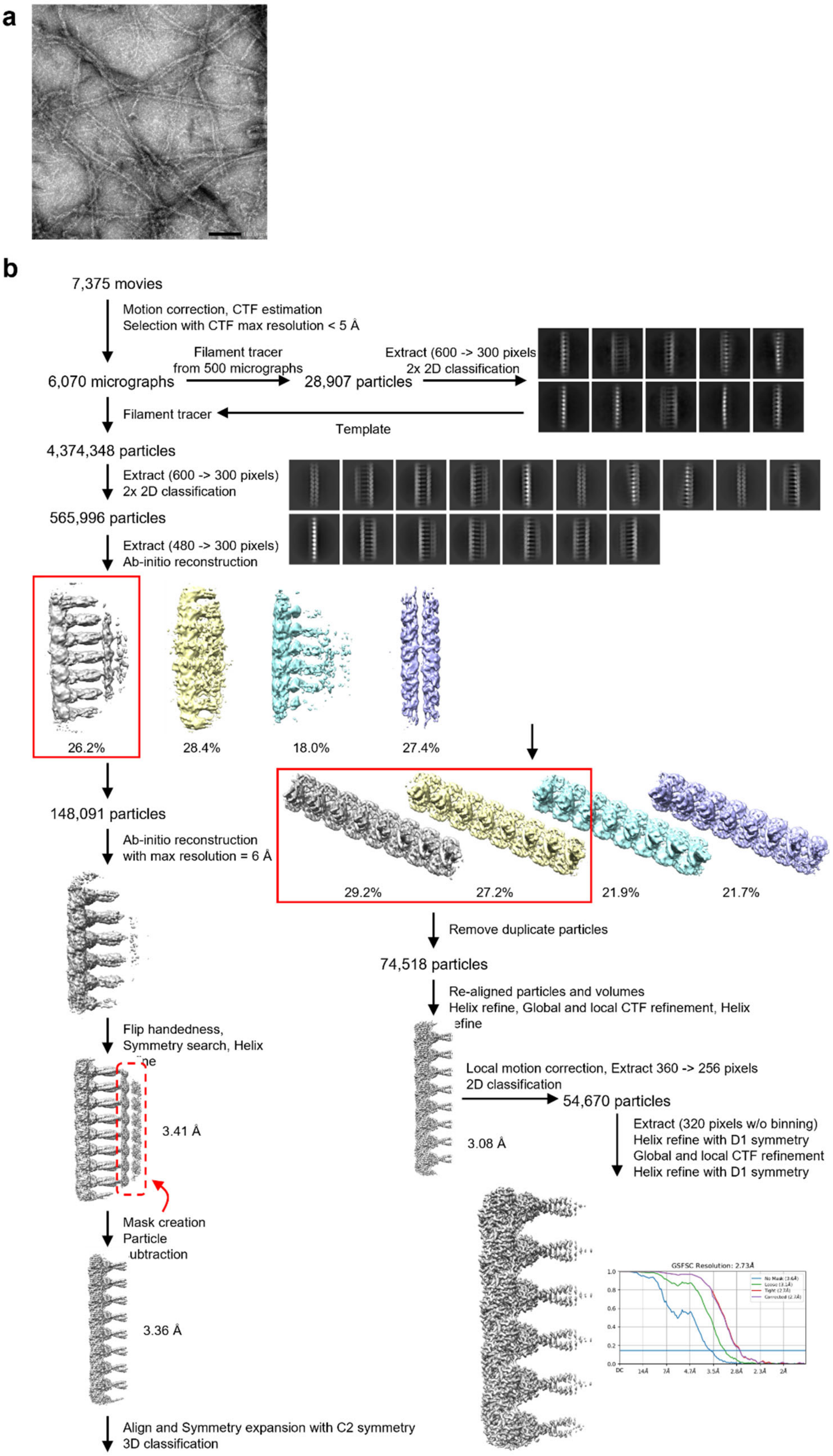

**Extended Data Fig. 1: Representative negative stain EM image and cryo-EM data processing workflow of the KpFtsZ-ZapA complex. a,** A negative stain EM image. The scale bar represents 100 nm. **b,** Cryo-EM data processing workflow of the KpFtsZ-ZapA complex.

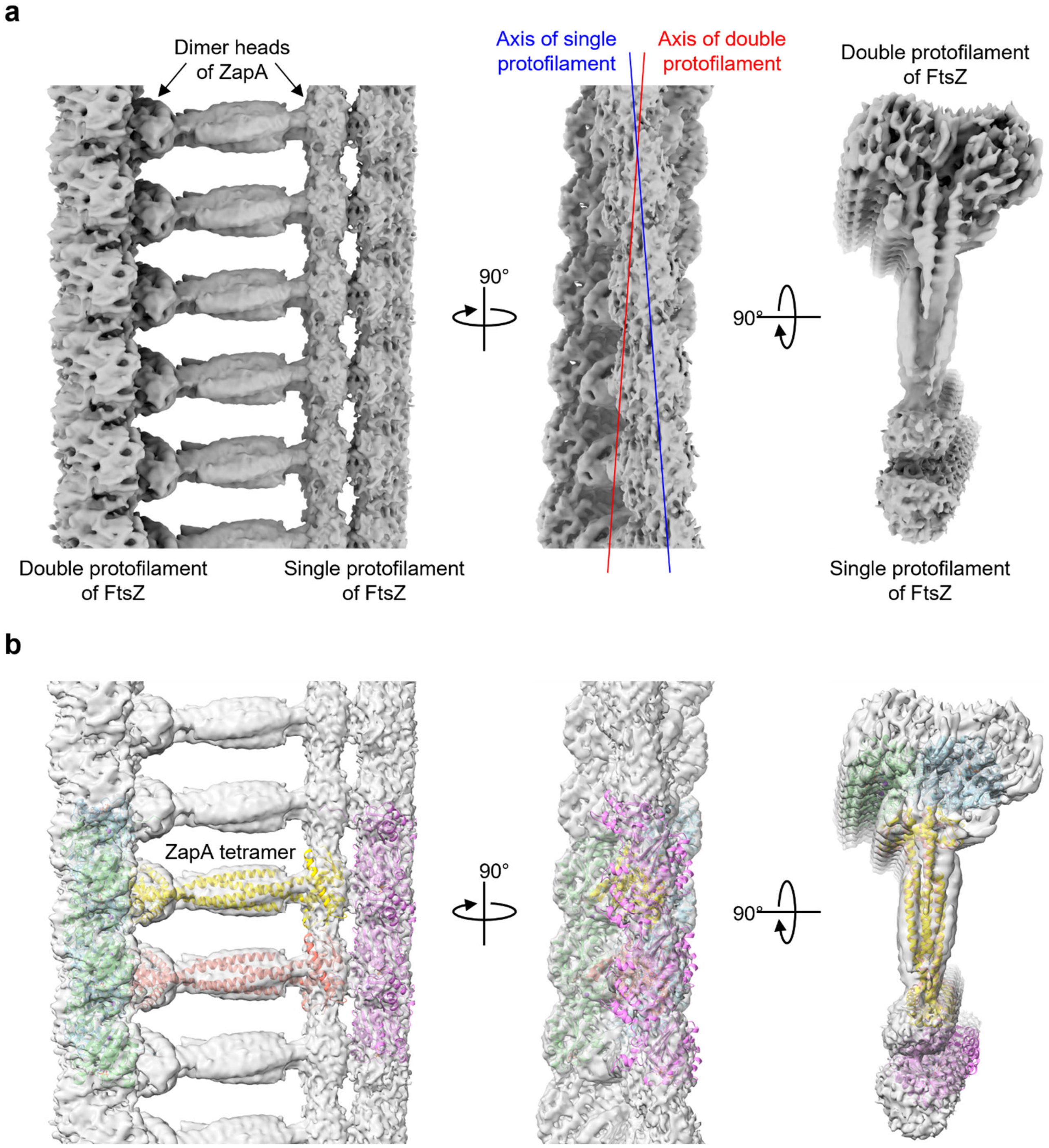

**Extended Data Fig. 2: Re-centered cryo-EM map of the ZapA-FtsZ complex. a,** Three orthogonal views of the map. Axes of FtsZ double and single protofilaments are also shown in the middle panel. **b,** Fitted FtsZ and ZapA models with the same map in a. Each of FtsZ protofilaments and ZapA tetramers is shown in different colors. Dimer head bound to the single protofilament is rotated from that bound to the double protofilament. The orientation of the single protofilament (shown in magenta) is not accurate.

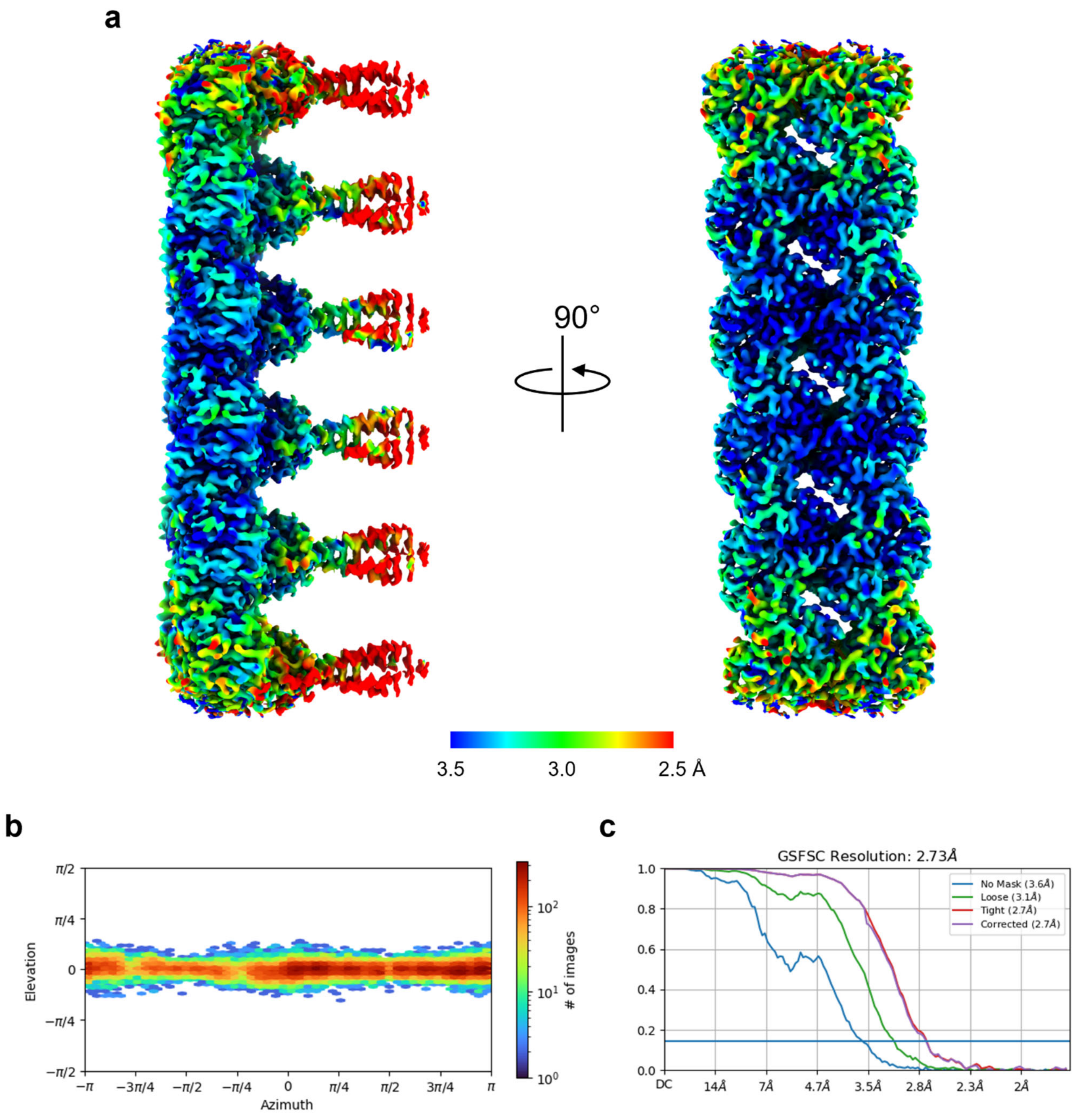

**Extended Data Fig. 3: Cryo-EM data processing of the ZapA-FtsZ complex. a**, Final sharpened map colored by local resolution as in the color bar. **b**, Angular distribution of the particles used in the final reconstruction. **c**, The FSC curve for the final map. The horizontal blue line indicates the FSC = 0.143 criterion.

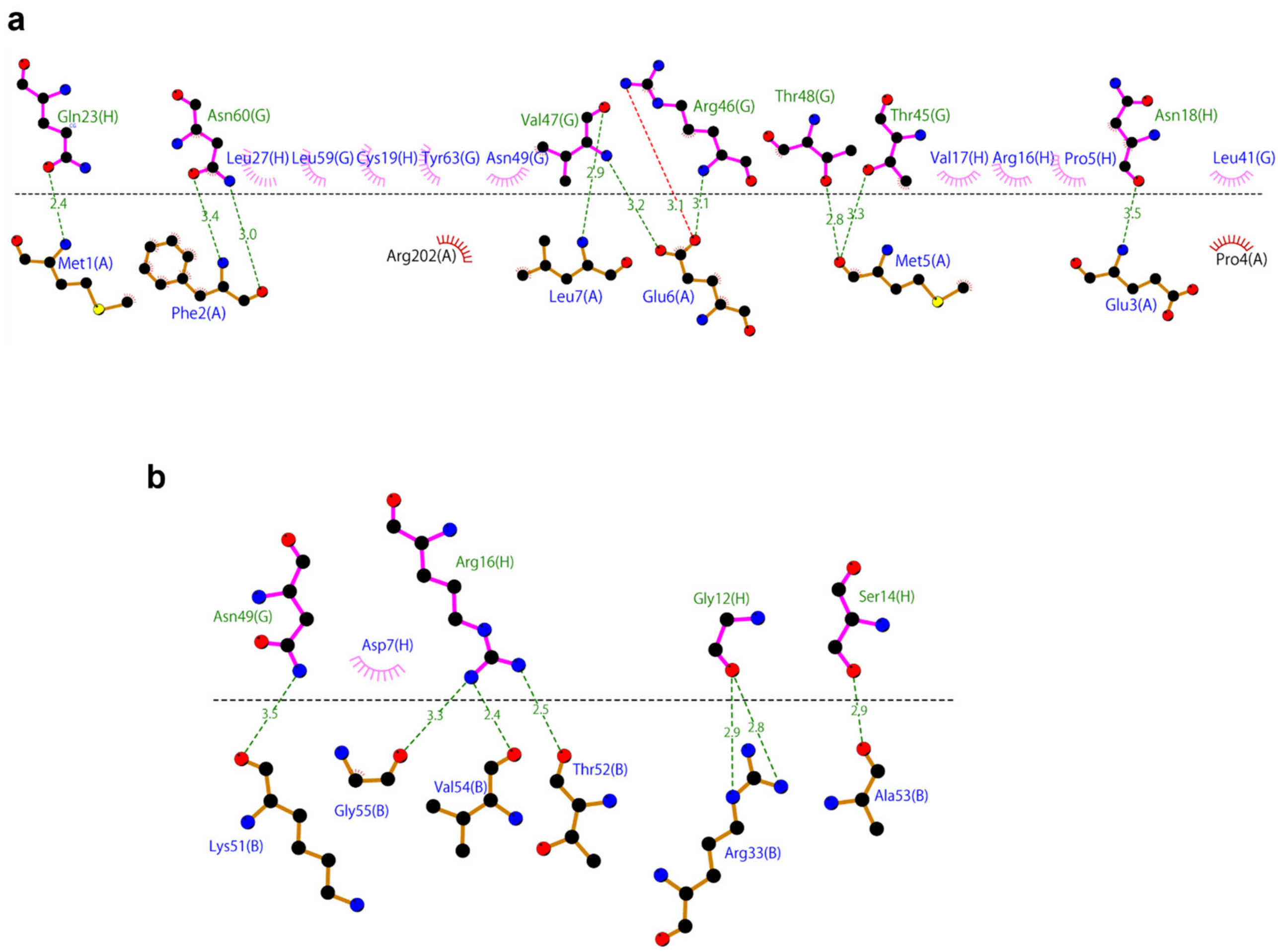

**Extended Data Fig. 4: Detailed description of the interactions generated with the program Ligplot. a**, Interactions between ZapA (chains G and H) and FtsZ (chain A). Hydrogen bonds are represented by dashed lines. **b**, Interactions between ZapA (chains G and H) and FtsZ (chain B).

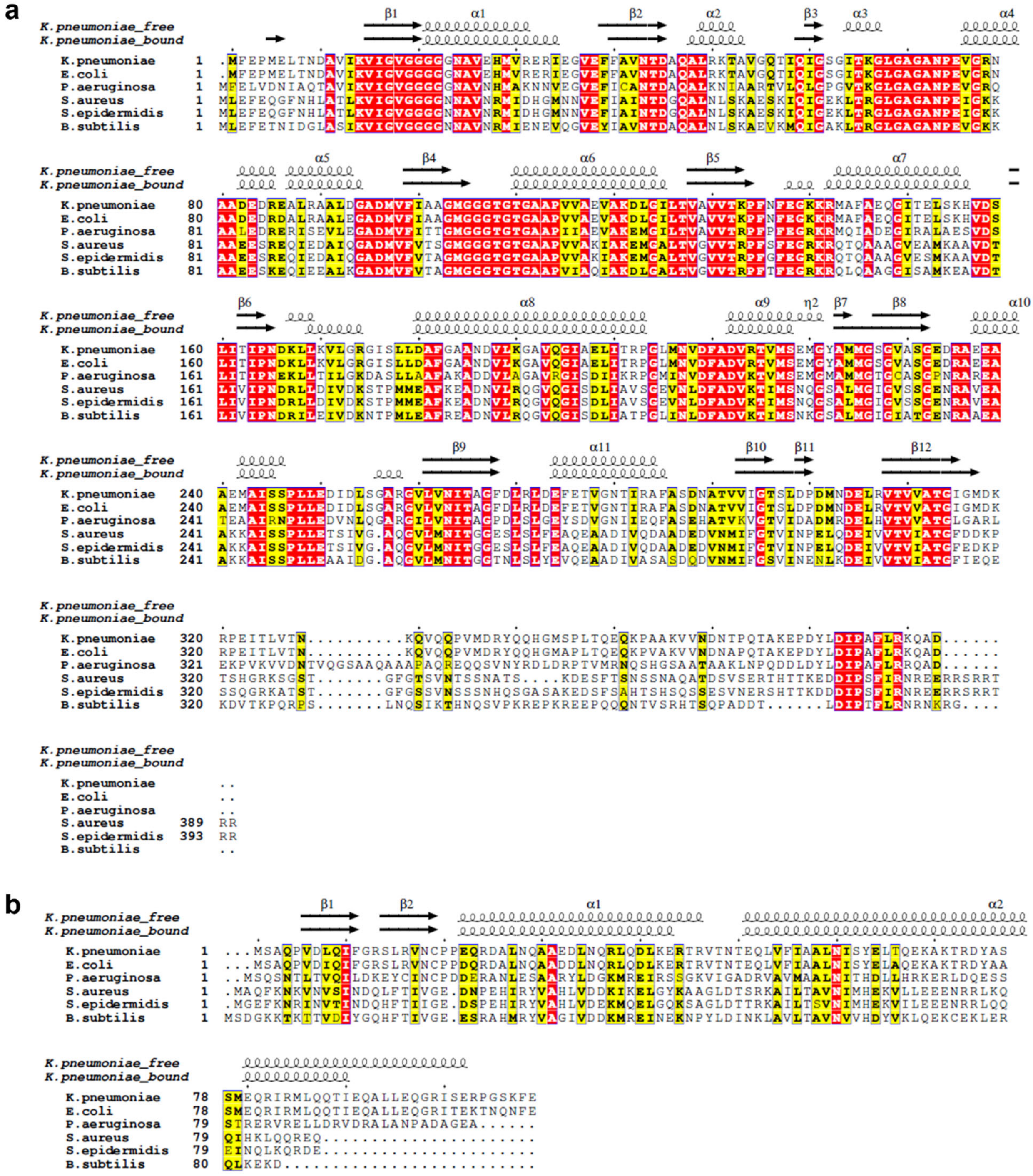

**Extended Data Fig. 5: Multiple sequence alignment of FtsZ and ZapA across species:** *K. pneumoniae* FtsZ (KpFtsZ; UniProt ID A6T4N8), *E. coli* FtsZ (EcFtsZ; UniProt ID P0A9A6), *P. aeruginosa* FtsZ (PaFtsZ; UniProt ID P47204), *B. subtilis* FtsZ (BsFtsZ; UniProt ID P17865), *S. aureus* FtsZ (SaFtsZ; UniProt ID Q6GHP9) *S. epidermidis* FtsZ (SeFtsZ; UniProt ID Q8CPK4), *K. pneumoniae* ZapA (KpZapA; UniProt ID B5XUC8), *E. coli* ZapA (EcZapA; UniProt ID P0ADS2), *P. aeruginosa* ZapA (PaZapA; UniProt ID Q9HTW3), *B. subtilis* ZapA (BsZapA; UniProt ID W8URK0), *S. aureus* ZapA (SaZapA; UniProt ID A0A380DNW5) and *S. epidermidis* ZapA (SeZapA; UniProt ID A0A0N1EE61).

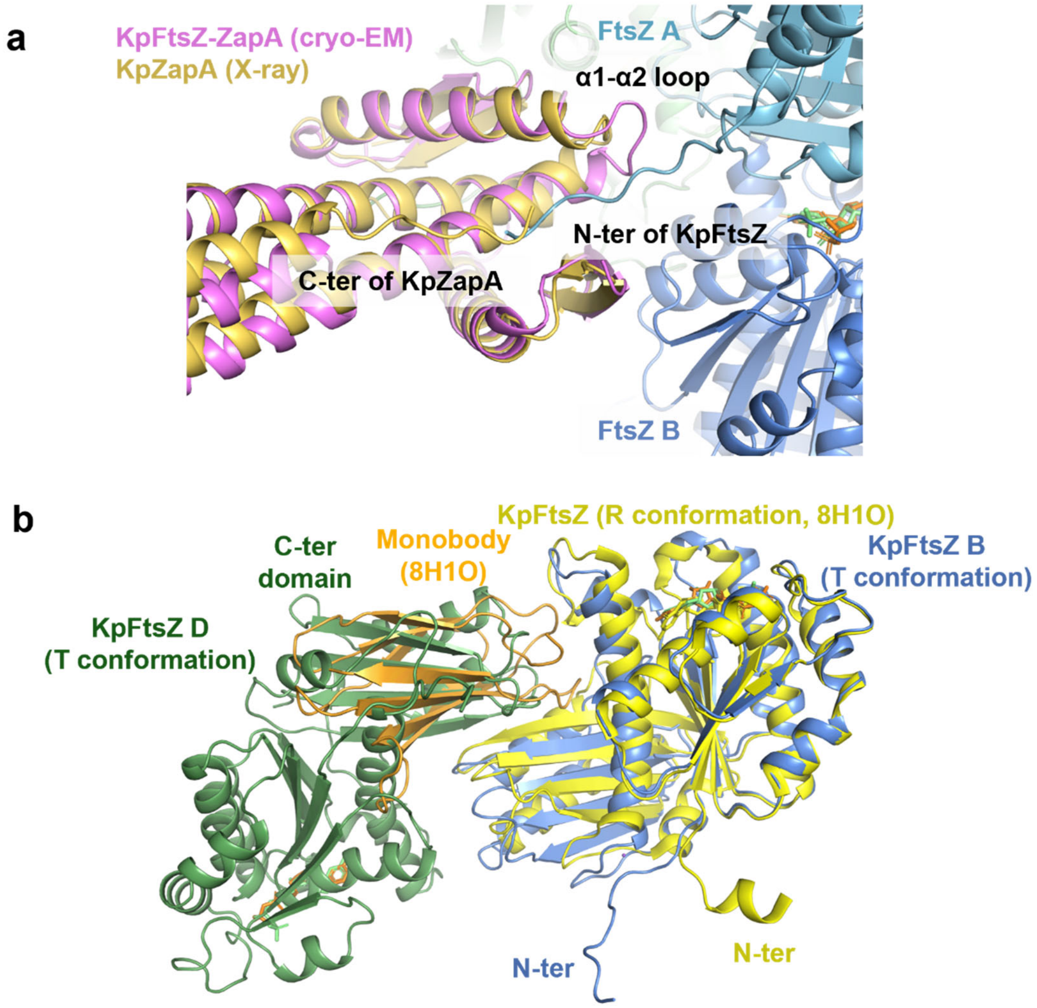

**Extended Data Fig. 6: Structural comparison of FtsZ within the ZapA-FtsZ complex. a,** Overlapping of N-terminus of KpFtsZ and C-terminus of KpZapA. The coloring is the same as in Fig. 2c. **b,** Overlapping of C-terminal domain in KpFtsZ from another protofilament and the monobody bound to KpFtsZ in the R conformation.

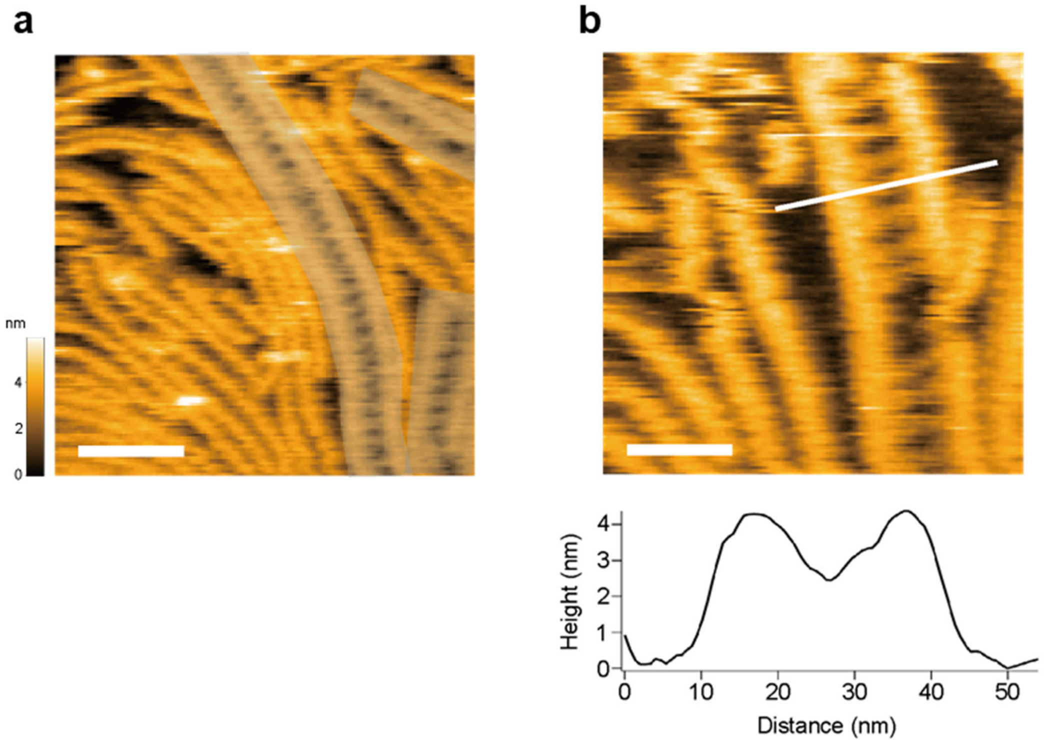

**Extended Data Fig. 7: HS-AFM Image of the ZapA-FtsZ Complex. a,** The shaded region shows a ladder-like structure where two single FtsZ filaments are cross-linked by ZapA tetramers. Scale bar: 50 nm, frame rate: 1 fps. **b,** Enlarged image of the ladder-like structure and its cross-sectional profile along the white line on the image. Scale bar: 25 nm, frame rate: 2 fps.

**Extended Data Table 1: CryoEM data collection and processing**

| Dataset | FtsZ double filament-ZapA tetramer |
|---|---|
| EMDB accession code | EMD- 60837 |
| PDB accession code | 9ISK |
| **Data collection and processing** | |
| Magnification | 60,000 |
| Voltage (kV) | 300 |
| Electron exposure ($e^{-}/Å^{2}$) | 60 |
| Defocus range (μm) | −0.5 to −2.0 |
| Pixel size (Å) | 0.878 |
| Symmetry imposed | D1 helical |
| Imported movies (no.) | 7,375 |
| Initial particle images (no.) | 4,374,348 |
| Final particle images (no.) | 54,670 |
| Map resolution (Å) | 2.73 |
| FSC threshold | 0.143 |
| Helical parameters | |
| Rise (Å) | 44.58 |
| Twist (°) | -3.11 |
| **Refinement** | |
| Initial model used (PDB code) | 8IBN, 4P1M |
| Model resolution (Å) | 2.6/2.7/3.0 |
| FSC threshold | 0/0.143/0.5 |
| Model vs. Data CC (mask) | 0.84 |
| (volume) | 0.82 |
| Model composition | |
| Non-hydrogen atoms | 17,586 |
| Protein residues | 2,324 |
| Ligands | 6 (GMPCPP) |
| | 6 (K) |
| R.m.s. deviations | |
| Bond lengths (Å) | 0.002 |
| Bond angles (°) | 0.536 |
| Validation | |
| MolProbity score | 1.49 |
| Clashscore | 7.90 |
| Rotamer outliers (%) | 0 |
| Ramachandran plot | |
| Favored (%) | 97.74 |
| Allowed (%) | 2.26 |
| Outliers (%) | 0 |

**Extended Data Table 2: Protein-protein interface analysis by PDBePISA server.**

| Chain 1:Chain 2 | Interface, Å$^2$ | $\Delta^{i}G$, kcal |
|---|---|---|
| ABEF:GH | 2105.4 | –17.0 |
| A:B | 1390.6 | –16.0 |
| B:EF | 523.4 | –3.6 |
| B:F | 207.7 | 1.0 |
| B:E | 315.7 | –4.6 |
| ABC:DEF | 1268.2 | –10.6 |

**Extended Data Table 3: Crystallographic data collection and refinement statistics.**

| Dataset | KpZapA |
|---|---|
| PDB accession code | 9ISJ |
| **Data collection and processing** | |
| Space group | $P6_122$ |
| Cell dimensions | |
| *a*, *b*, *c* (Å) | 54.173, 54.173, 330.044 |
| α, β, γ (°) | 90.00, 90.00, 120.00 |
| Total reflections | 250,986(22,250)* |
| Unique reflections | 28,055(2,712) |
| Resolution range | 46.45–1.80(1.86–1.80) |
| $R_{merge}$ | 0.05109(1.101) |
| $I/\sigma I$ | 20.03(1.66) |
| Completeness (%) | 99.84(100.0) |
| Redundancy | 9.0(8.2) |
| $CC_{1/2}$ | 0.999(0.872) |
| **Refinement** | |
| $R_{work}/R_{free}$ | 0.215/0.239 |
| No. of atoms | |
| Protein | 1712 |
| Ligand | 1 |
| Water | 134 |
| *B* factors ($Å^2$) | |
| Protein | 45.74 |
| Ligand | 57.39 |
| Water | 48.94 |
| R. m. s. deviations | |
| Bond length (Å) | 0.0089 |
| Bond angles (°) | 1.6067 |
| Validation | |
| MolProbity score | 0.95 |
| Clashscore | 1.75 |
| Rotamer outliers (%) | 1.06 |
| Ramachandran plot | |
| Favored (%) | 100.00 |
| Allowed (%) | 0.00 |
| Outliers (%) | 0.00 |

*Each dataset was collected from one crystal. *Values in parentheses are for highest-resolution shell.

Movie 1: HS-AFM movie demonstrating the dynamic interaction between ZapA and an FtsZ filament.

Frame rate: 2 fps, 3× playback speed.
